# LCA robustly reveals subtle diversity in large-scale single-cell RNA-seq data

**DOI:** 10.1101/305581

**Authors:** Changde Cheng, John Easton, Celeste Rosencrance, Yan Li, Bensheng Ju, Justin Williams, Heather L Mulder, Wenan Chen, Xiang Chen

## Abstract

Single-cell RNA sequencing has emerged as a powerful tool for characterizing the cell-to-cell variation and dynamics. We present Latent Cellular Analysis (LCA), a machine learning– based analytical pipeline that features a dual-space model search with inference of latent cellular states, control of technical variations, cosine similarity measurement, and spectral clustering. LCA has proved to be robust, accurate, scalable, and powerful in revealing subtle diversity in cell populations.

## Backgrounds

Single-cell RNA sequencing (scRNA-seq) enables the characterization of cell-to-cell variation in transcript abundance, leading to a deep understanding of the diversity of cell types and the dynamics of cell states in populations of thousands to tens of thousands of single cells [1-3]. Although scRNA-seq offers enormous opportunities and has inspired a tremendous explosion of data-analysis methods for identifying heterogeneous subpopulations, significant challenges can arise because of the inherently high noise associated with data sparsity and the ever-increasing number of cells sequenced. The current state-of-the-art algorithms have significant limitations. The biological meaning of the cell-to-cell similarity learned by most machine learning–based tools (such as Seurat [4], Monocle2 [5], SIMLR [6], and SC3 [7]) is difficult to interpret. Several methods require the user to provide an estimation of the number of clusters in the data, and this may not be readily available. Furthermore, most methods have a high computational cost that will be prohibitive for datasets representing large numbers of cells. Lastly, although certain technical biases (e.g., cell-specific library complexity) have been recognized as major confounding factors in scRNA-seq analyses [8], other technical variations (e.g., batch effects and systematic technical variations that are irrelevant to the biological hypothesis being evaluated) have received minimal attention, even though they present major challenges to the analyses [9]. Most methods employ a gene-selection step before clustering analysis, based on the assumption that a small subset of highly variable genes is most informative for revealing cellular diversity. Although this assumption may be valid in many scenarios, it fails when informative genes are not most variable. This can happen when the difference among subpopulations is subtle, or there is a strong batch effect, while most variable genes differ by batch.

To address these issues, we developed Latent Cellular Analysis (LCA), an accurate, robust, and scalable computational pipeline that will facilitate a deep understanding of the transcriptomic states and dynamics of single cells in large-scale scRNA-seq datasets. LCA makes a robust inference of the number of populations directly from the data (a user can specify this with a priori information), rigorously models the contributions from potentially confounding factors, generates a biologically interpretable characterization of the cellular states, and better reveals the underlying population structures. Furthermore, LCA addresses the scalability problem by learning a model from a subset of the sample, after which a theoretical scheme is used to assign the remaining cells to identified populations.

## Results

### Overview of the LCA method

LCA employs a heuristic dual-space search approach (**Fig. 1**) to provide an optimal model for subpopulation structure identification (including the removal of confounding factors, the inference of the number of clusters and informative states, and a mapping function from the expression vector to cluster membership). In the primary latent cellular (LC) space, LCA bypasses gene selection and performs LC state inference from the global gene expression matrix, removes cellular states representing known confounding factors (e.g., cell-specific library complexity and batch information), and measures cell-4 to-cell similarity by using the cosine of the angle between the low-dimensional cellular-state vectors. Spectral clustering is employed to derive a set of candidate clustering models with a range of cluster numbers. Meanwhile, in the dual principle component (PC) space, cells are projected into a low-dimensional space based on principal component analysis (PCA) of the expression vector. The list of significant components is determined by the Tracy-Widom test [10], and cell-to-cell similarity is measured by correlation similarity in the PC space. LCA then ranks candidate clustering models derived in the LC space by the silhouette index [11] measured in the PC space. Finally, LCA retains informative cellular states from the selected clustering model(s) and uses these states to update the final clustering solution(s). Although the "optimal model" is not necessarily the solution with the best numerical score, it should be among the top-ranked models. Therefore, LCA provides users with multiple top-ranked solutions for biology-based evaluations.

**figure 1.**
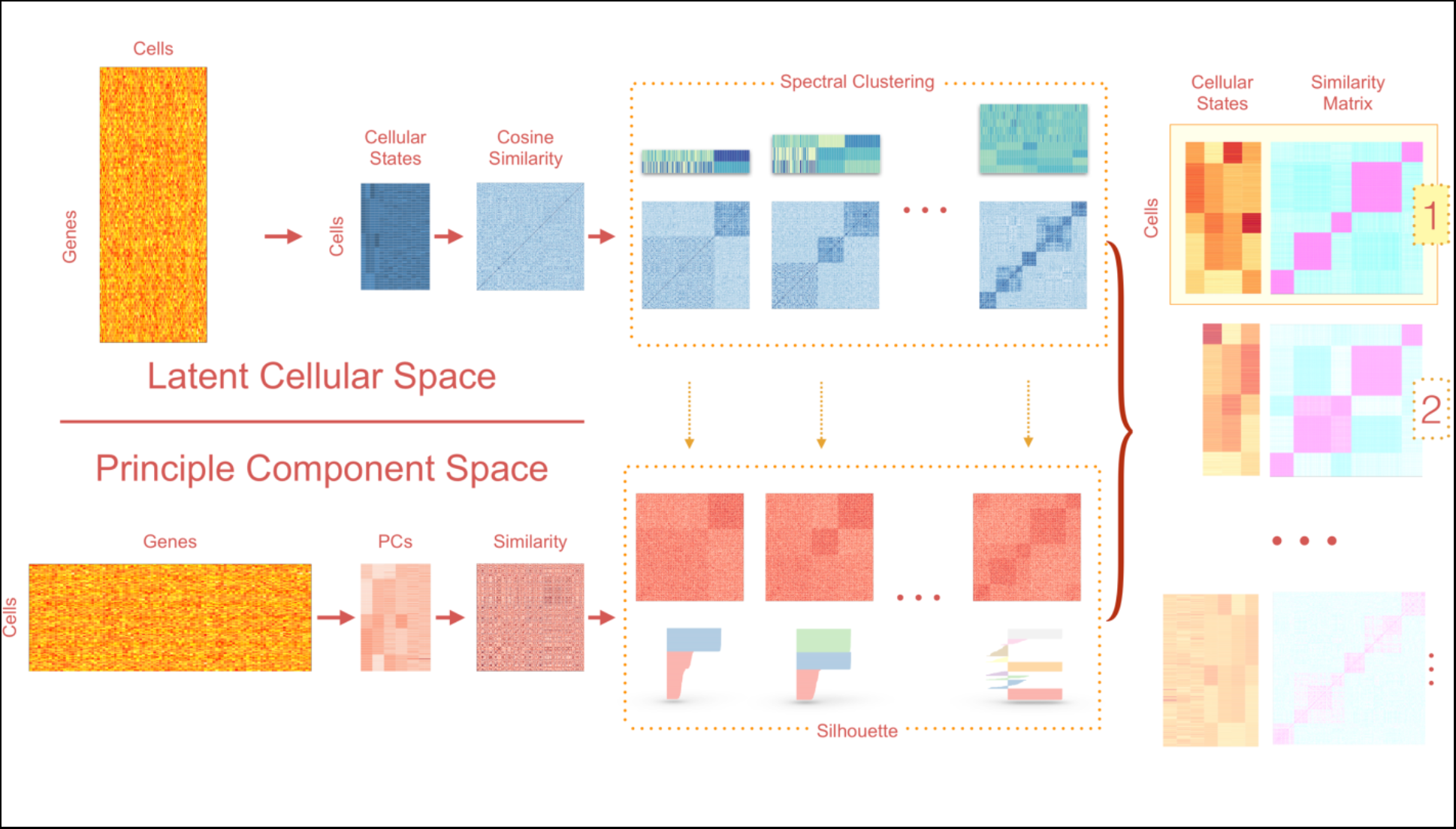
Overview of the workflow of LCA. LCA infers LC states from full expression matrices. Explicit gene filtering is not necessary. LCA converts the raw transcript count data to gene fractions and then performs log transformations. The algorithm features a dual-space model search, generating candidate clustering models based on the cosine similarity matrix in the LC space. Candidate models are then ranked based on the silhouettes measured in the PC space.

### LCA accurately identifies cellular states in simulated datasets

We benchmarked LCA with four commonly used state-of-the-art scRNA-seq clustering algorithms (SC3 [7], Seurat [4], Monocle2 [5], and SIMLR [6]) in 3000 simulated datasets. LCA outperformed competing methods in subpopulation number inference, cluster membership inference, and running time.

We simulated three equally spaced cell subpopulations of various sizes by shuffling a predetermined number of genes across cells from high-quality, Unique Molecular Identifier (UMI)-based scRNA-seq data. We simulated seven scenarios with increasing signal strength, varying the number of simulated differentially expressed genes (DEGs), the fold change among groups, and the average expression levels of the simulated DEGs. We also investigated the effect of the sample size (from 250 to 4000 cells) on the performance of the algorithms.

A typical clustering analysis partitions data points into meaningful subgroups without a priori knowledge of the number of subsets or their compositions. An essential component of clustering analysis is inferring the number of clusters from the data [12], which presents an analytical challenge distinct from that posed by clustering with a given number of clusters. Therefore, we first analyzed the accuracy and robustness of the subpopulation number inference. Monocle2 and SIMLR were excluded from this analysis because both algorithms require an input of the number of clusters from end-users. **Fig. 2A** shows that although LCA started with a slightly off-target estimation of the cluster number in datasets with very weak signal strength, it stably converged to the correct number with increasing signal strength from the number of cells, the number of DEGs, the fold difference, or the expression level. In contrast, using SC3 or Seurat resulted in an upward bias in the estimation of the number of clusters, and this bias increased with both the sample size and the signal strength. For example, in simulated model #6, with the strongest signal strength, the actual cluster number remained the same (three) for all sample sizes, but SC3 estimated medians of four, eight, and sixteen clusters in datasets with 250, 1000, and 4000 cells, respectively. Seurat estimated medians of three, four, and nine subpopulations in the same respective datasets. Combining the results of all simulations, we find that LCA arrived at the correct number of clusters in 70.2% of simulations, which is a significantly higher percentage than was achieved with SC3 (5.6%, *P*< 2.2 × 10^−16^ [proportion test]) and Seurat (22.7%, *P*< 2.2 × 10^−16^ [proportion test]).

**figure 2.**
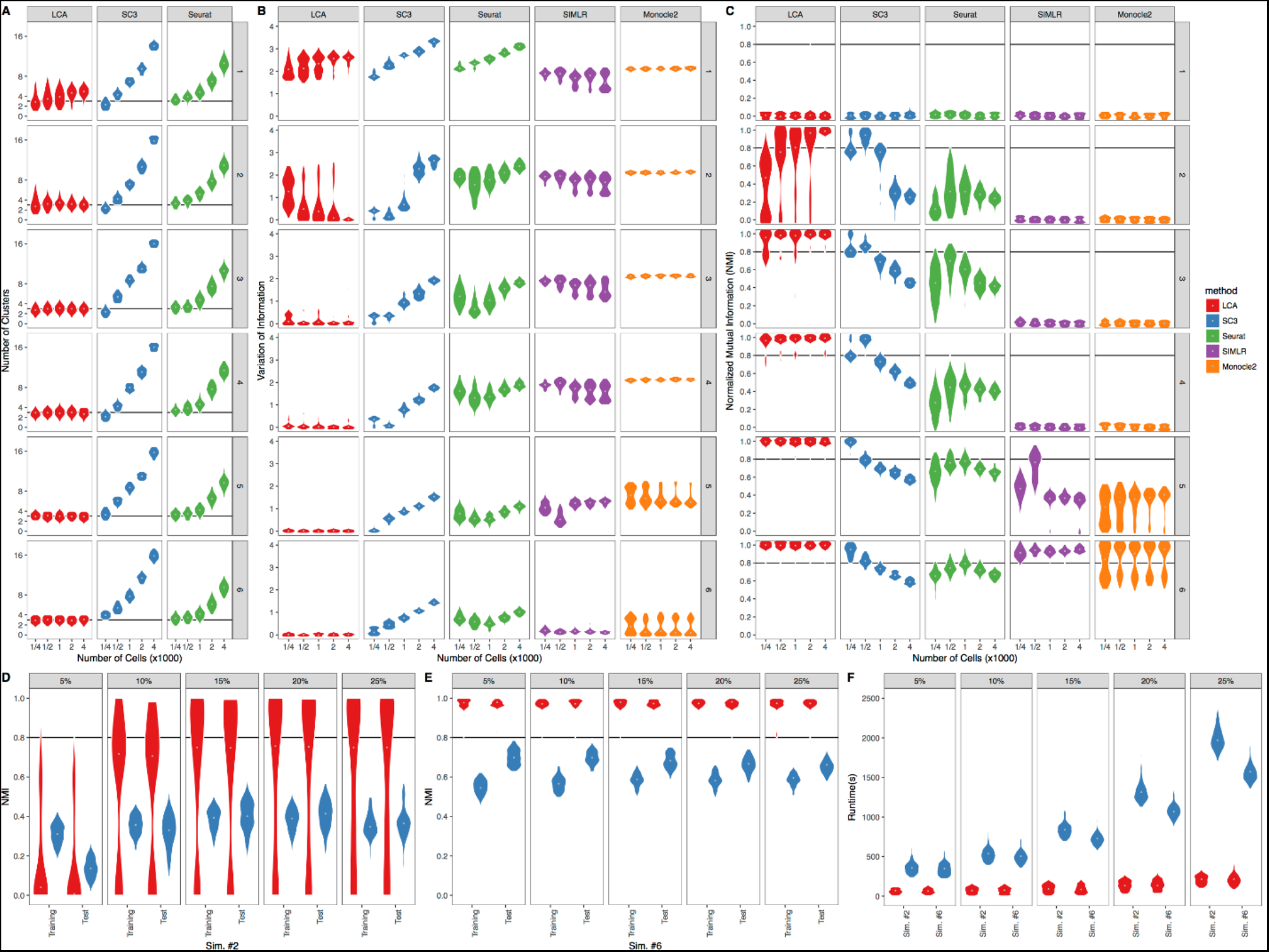
Benchmarking of LCA against four other methods with simulated datasets. (A) The number of clusters inferred by LCA, SC3, and Seurat was compared for seven simulated datasets: the signal strength increases from simulated datasets 1 to 6. Each simulated dataset contained 100 randomly generated samples. (B) Comparison of the clustering performance of LCA, SC3, Seurat, Monocle2, and SIMLR, as measured by variation of information, across the seven simulated datasets. For LCA, SC3, and Seurat, the number of clusters was inferred during the analysis. For Monocle2 and SIMLR, we used the correct number of clusters. (C) Comparison of the clustering performance as measured by normalized mutual information. (D) Clustering performance scalability of LCA and SC3. Each panel represents the results obtained with a different percentage of cells used for training (from 5% to 25%). The total number of cells in the sample was 4000. The signal in this comparison was relatively weak. (E) Comparison of the clustering performance scalability of LCA and SC3 when the signal was strong. (F) Comparison of the running times of LCA and SC3.

We used the variation of information (VI [13], 0 for perfect matching) (**Fig. 2B**) and normalized mutual information (NMI [14], 1 for perfect matching) (**Fig. 2C**) to measure the clustering accuracy. For methods with a feature for inferring the number of clusters from the data (LCA, SC3, and Seurat), we compared the ground-truth labels with inferred cluster membership from the optimal clustering models. For methods without this inference feature (Monocle2 and SIMLR), the optimal models were derived with the correct number of clusters (three). The comparison showed that LCA outperformed other methods in most simulations (**Fig. 2**). When all 3000 simulation datasets were combined, LCA achieved the highest overall accuracy (average NMI = 0.756; VI = 0.555). Specifically, when LCA correctly inferred the number of clusters (in 70.2% of all simulations), it produced near-perfect accuracy (average NMI = 0.941, VI = 0.122). The accuracies of the competing algorithms were consistently lower. Notably, SC3 achieved a relatively good performance (NMI = 0.807, VI = 0.389) when it happened to find the correct number of clusters (mostly in small datasets). However, its performance deteriorated for other datasets, primarily due to an overestimation of cluster numbers. The overall accuracy of SC3 was significantly lower than that of LCA (NMI = 0.583, *P* < 2.2 × 10^−16^; VI = 1.203, *P* < 2.2 × 10^−16^ [Wilcoxon signed-rank test]). More strikingly, even if provided with the correct cluster number, Monocle2 attained high accuracy only for the dataset with the strongest signal strength (simulated model #6), and its performance was suboptimal compared to that of LCA (NMI = 0.190, *P* < 2.2 × 10^−16^; VI = 1.719, *P* < 2.2 × 10-^16^ [Wilcoxon signed-rank test]). Similarly, the overall accuracies of Seurat (NMI = 0.415, *P* <2.2 × 10^−16^; VI = 1.545, *P* < 2.2 × 10^−16^ [Wilcoxon signed-rank test]) and SIMLR (NMI = 0.231, *P* < 2.2 × 10^−16^; VI = 1.353, *P* < 2.2 × 10^−16^ [Wilcoxon signed-rank test]) were significantly lower than that of LCA.

### Scalability to large data sets

In addition to the superior accuracy of LCA with respect to both cluster number and membership assignment, its implementation renders exceptional scalability, which is an attractive feature for scRNA-seq analysis of an ever-increasing number of cells. LCA can derive a cell subpopulation structure by using a relatively small representative set (training cells) sampled from the full data. LCA provides mathematical formulae with which to project the remaining cells (testing cells) directly to the inferred low-dimensional LC space, after which individual cells are assigned to the subpopulation with the best similarity. Consequently, LCA runs at a low level of computational complexity. The median overall running times for a 4000-cell dataset with 10% or 25% training cells were 40.6 s and 202.9 s, respectively. We compared the accuracy of clustering between the training cells (5% – 25% of all cells) and the remaining testing cells in two simulated models (**Fig. 2C, D**, models #2 and #6). As expected, the accuracy of the clustering model improved with the increase of training cells in the simulated dataset #2 (weak signal strength) and remained at a near-perfect level in simulated dataset #6 (strong signal strength). Strikingly, LCA achieved comparable accuracies in the testing cells in both models and outperformed SC3 in terms of accuracy and running time.

### Evaluation of LCA on published data sets

By using publicly available large-scale scRNA-seq datasets, we demonstrated that LCA identified orthogonal LC states of biological importance and produced parsimonious and accurate models that were highly consistent with biological knowledge.

We evaluated the performance of LCA with large-scale datasets published by Zheng et al. [15] and Tirosh et al. [16]. Using the GemCode platform (10x Genomics, Pleasanton, CA), Zheng et al. generated reference scRNA-seq transcriptomes for subpopulations of peripheral blood mononuclear cells (PBMCs) purified via well-established cell surface markers [15]. We generated a large-scale purified T-cell dataset (representing 55,000 cells) by combining the CD4+ T-helper (CD4+ helper), CD4+/CD25+ regulatory T (T_reg_), CD4+/CD45RO+ memory T (T_mem_), CD4+/CD45RA+/CD25- naïve T (CD4+ ab T), CD8+ cytotoxic T (cytotoxic T), and CD8+/CD45RA+ naïve cytotoxic T (CD8+ ab T) cell subpopulations. We inferred the cellular states and subpopulation structure by using 10% and 25% of the full dataset, and we assigned the remaining cells to one of the inferred subpopulations. We evaluated the top three models for both runs, which were the 3-population, 4-population, and 5-population models. Despite a 9.6-fold difference in running time, both the 3-population and 5-population models from the two runs achieved high consistency (NMI = 0.87 and 0.84, respectively). Although the 4-population models differed between runs, they represented two different subpopulation-merging orders from the 5-population model to the 3-population model. We selected the 5-population model learned from 25% cells (with 19 LC states) for further biological inference. Whereas the purified T_mem_, CD4+ ab T, and CD8+ ab T cells contained cells mostly from a single subpopulation (Clusters 1, 3, and 4, respectively), different levels of heterogeneity were detected in the remaining purified populations, especially the CD4+ helper and cytotoxic T cells. Given the single surface-marker settings in the purification of CD4+ helper and cytotoxic T cells, it is not surprising that we found substantial heterogeneity in the two populations. Nevertheless, LCA inferred a parsimonious subpopulation structure in this large dataset (compared to the 1108 subpopulations predicted by SC3). We selected representative genes encoding surface markers, transcription factors, and secreted effector molecules for 19 usual T-cell subsets [17] and derived a PCA projection of the six purified populations and five inferred clusters based on the average population/cluster expression level of individual marker genes. The first PC largely described the PC represented the difference between the CD4+ subsets and CD8+ subsets. As expected, Clusters 3 and 4 were found next to the CD8+ ab T and CD4+ ab T cells. Cluster 1 was found near the T_mem_ population. Cluster 2 consisted mostly of T_reg_ cells and a smaller fraction of CD4+ helper cells that was located adjacent to the T_reg_ population. Both the CD4+ helper and cytotoxic T-cell populations were split approximately equally between naïand differentiated cells and were spotted between their corresponding naïve clusters (Cluster 3 for CD8+ cells and Cluster 4 for CD4+ cells) and differentiated clusters (Clusters 1 and 2 for CD4+ cells and Cluster 5 for CD8+ cells). Analysis of selected genes validated the separation of naïve and differentiated cells in the CD4+ helper and cytotoxic T cells (**Fig. 3**). An evaluation of the first LC state inferred from the full expression data revealed a striking approximation to the expected distribution of naïve and differentiated cells in both purified populations and inferred clusters. Similarly, the second LC state recaptured the CD4+ and CD8+ difference in the dataset. These results showed that LCA identified LC states of biological importance and consequently produced a parsimonious and accurate model that was highly consistent with biological knowledge.

**figure 3.**
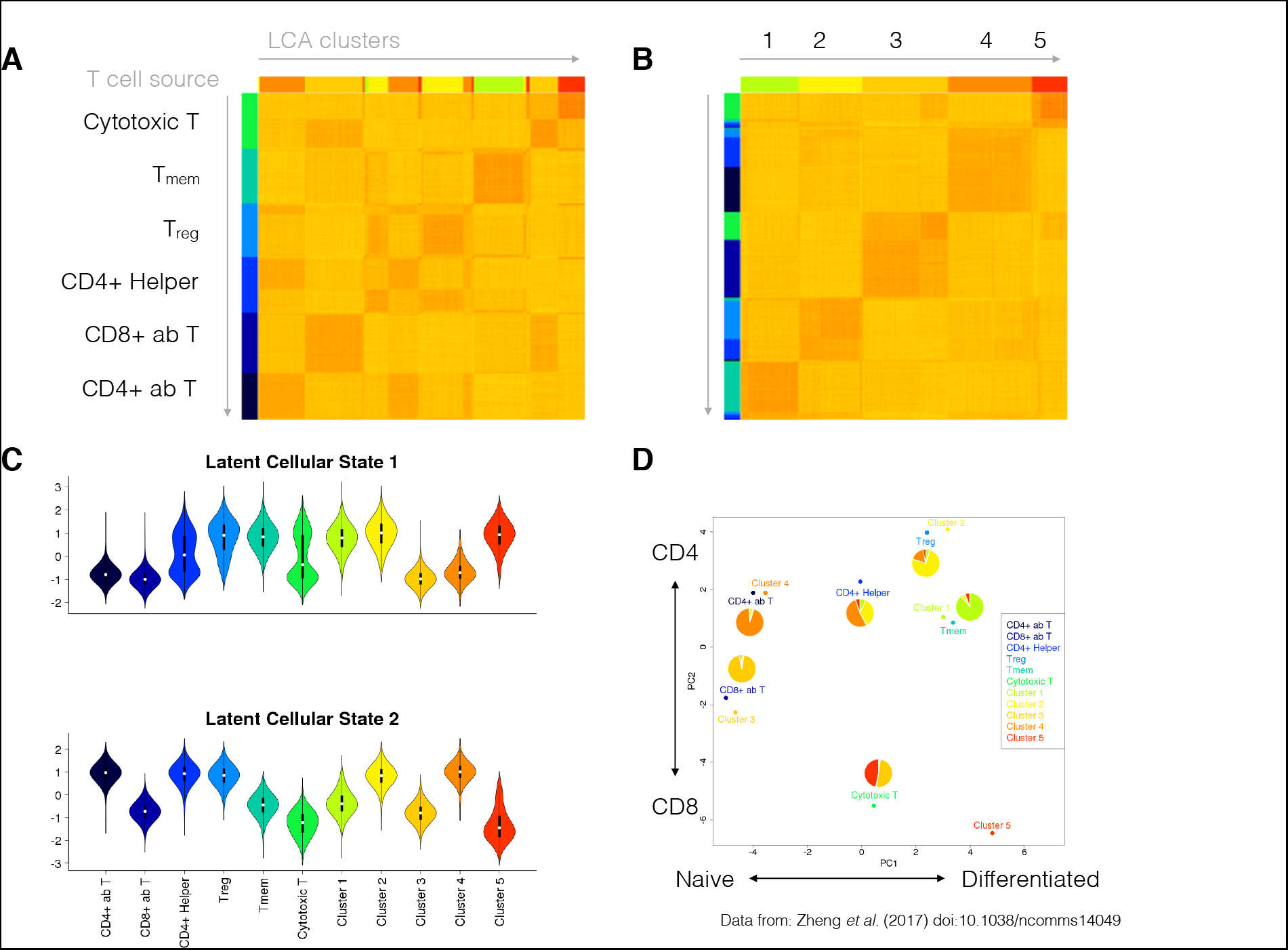
Investigation of PBMC heterogeneity by using LCA. We applied LCA to data from Zheng et al.^25^ for 55,000 cells representing a combination of six sorted T-cell populations. LCA identified five cell populations with biologically meaningful LC states. (A) Cell distance matrix, sorted by source ID. (B). Cell distance matrix, sorted by LCA clustering results. (C) The first LC state captured the difference between the naïve and differentiated T cells, whereas the second LC state captured the difference between the CD4+ and CD8+ cells. (D) LCA further revealed heterogeneity in the CD4+ helper T cells and CD8+ cytotoxic T cells.

We evaluated the performance of LCA in a second dataset published by Tirosh et al. [16], who employed a stepwise approach to analyze separately the malignant and stromal cells of 19 melanoma tumors captured on the C1 Fluidigm platform. We applied those authors’ cell-selection criteria [16] to generate a dataset with 1169 malignant cells from eight tumors and 2588 nonmalignant (stromal) cells. LCA inferred 18 clusters with 53 LC states (**Fig. 4**). Malignant cells dominated eight clusters, which were further separated by the patient origin of the tumors (**Fig. 4A**). Among the stromal cells, distinct clusters were identified for B cells, macrophages, cancer-associated fibroblasts, and endothelial cells (**Fig. 4B**). Moreover, LCA divided tumor-infiltrating T cells and natural killer cells into six clusters, which was concordant with the supervised analysis of T cells based on surface markers by Tirosh et al. [16]. Using the pre-defined marker genes for T-cell subsets and *MKI67* for cell-cycle activities, we revealed characteristics of these T-cell populations (**Fig. 4C**). Cluster 1 was enriched for naïve CD4+ cells, whereas Cluster 2 harbored mostly Treg and Tfh cells. Although Clusters 3 and 5 both had enriched signatures for cytotoxic T and exhaustive T cells, Cluster 3 had greater signal strength in both signatures, consistent with the reported high correlation between exhaustion markers and cytotoxic markers [16]. Cells in active cell cycles were grouped in Cluster 4, which showed unique enrichment of *MKI67* expression. Lastly, natural killer cells and T cells with weak cytotoxic activity and no exhaustion signatures were found in Cluster 6. As a comparison, we applied SC3 to the same data, which resulted in an estimation of 43 clusters. Although SC3 clustering of malignant cells was consistent with the separation by patient, accuracy in the stromal component was lower than with LCA (**Supplementary Table 1**). These results demonstrate the power of LCA to reveal subtle diversities in different subpopulations of tumor-infiltrating T cells in the presence of strong transcriptomic variations among malignant cells (from different patients) and various stromal cells from different lineages (cancer-associated fibroblasts, macrophages, B cells, and endothelial cells).

**figure 4.**
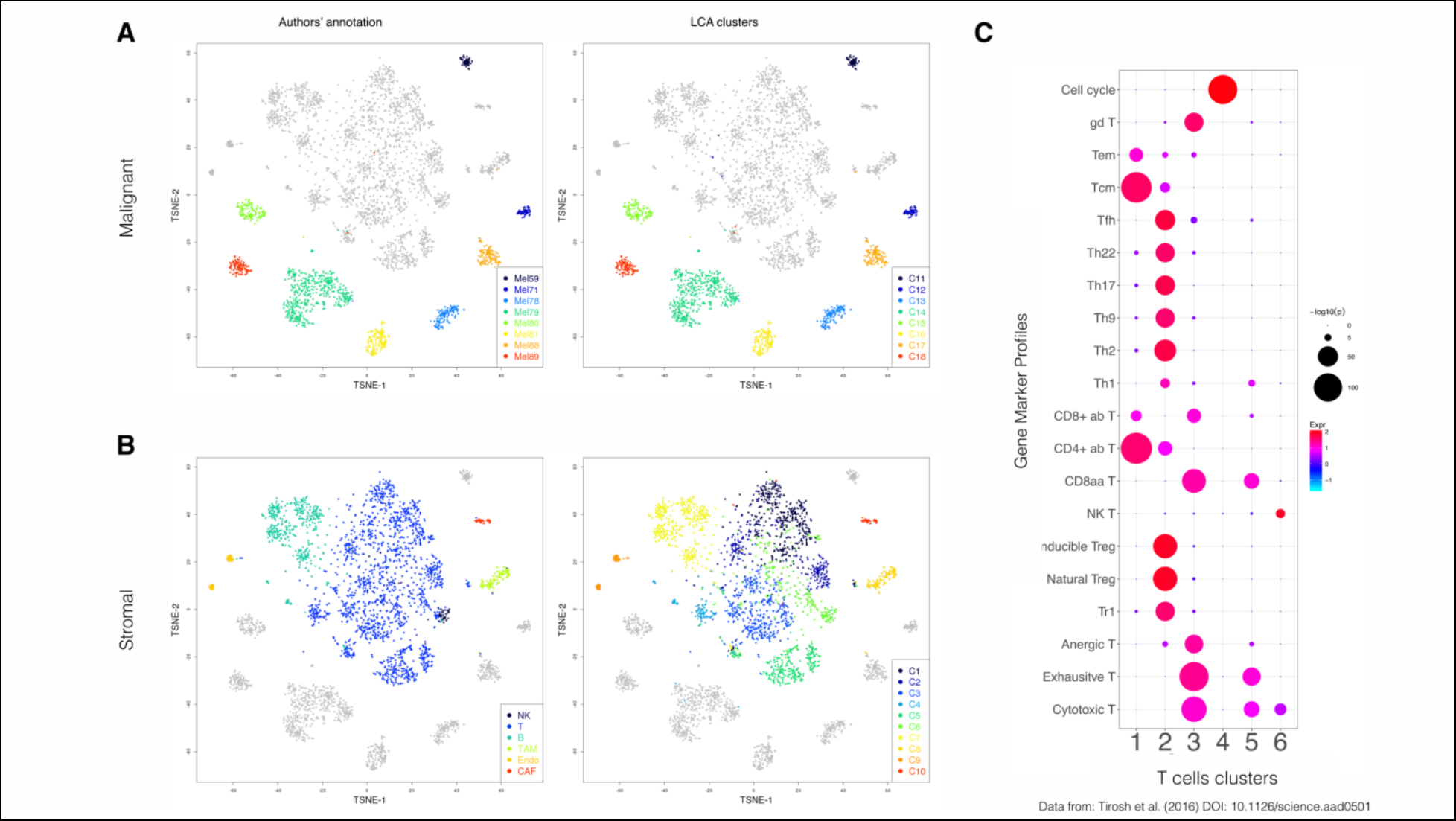
Reanalysis of melanoma cellular data [16] with LCA. We applied LCA to 1169 malignant cells from eight tumors and to 2588 stromal cells. LCA identified 18 clusters with a clear biological interpretation. LCA further revealed subtle diversity among the infiltrating T cells. (A) t-SNE plot of the clustering results for malignant cells, contrasting the LCA results and original results. (B) t-SNE plot of the clustering results for stromal cells, contrasting the LCA results and original results. (C) Enrichment of gene markers in T-cell clusters.

### LCA work robustly on a data set with strong batch effects

LCA groups cells based on orthogonal LC states aligned with major differences among cells, which enables control of technical variations (e.g., batch effects) without an explicit need of gene filtering. We evaluated the efficiency of batch effect removal in a cancer cell dataset and further experimentally validated the subpopulations revealed by LCA. Rhabdomyosarcoma is the most common soft-tissue tumor in children and has two major histologic subtypes with different genomic landscapes: *PAX3/PAX7-FOXO1* fusion–positive alveolar rhabdomyosarcoma (FP-ARMS) and fusion-negative embryonal rhabdomyosarcoma (ERMS) [18]. LCA identified two subpopulations in a scRNA-seq dataset for Rh41 cells (a commonly used human *PAX3-FOXO1* FP-ARMS cell line). The *CD44* gene, which encodes a commonly used cell surface marker with great prognostic and therapeutic potential [19, 20], appeared at the top of the differentially expressed genes (DEGs) of the two subpopulations (**Supplementary Figure 1A**). Flow cytometry confirmed a bimodal expression pattern of CD44 in Rh41 cells (**Supplementary Fig. 1B**). We first used bulk RNA-seq to profile unsorted populations and CD44^high^ and CD44^low^ subpopulations sorted by fluorescence-activated cell sorting. In addition to the differences among the sorted CD44^high^, CD44^low^, and unsorted populations, the analysis revealed strong batch effects. Specifically, samples in Batch 1 and those in Batches 2 and 3 were separated on the first PC, whereas the biologically different populations (the unsorted population and sorted subpopulations) were separated on the second PC (**Supplementary Fig. 1C**). We collected scRNA-seq data for the unsorted populations in Batch 1 and the sorted CD44^high^ and CD44^low^ subpopulations in Batch 2 (10x Genomics, Pleasanton, CA). The libraries were sequenced at different depths for the three samples. The median numbers of UMIs captured per cell were 8278, 6678, and 12,850 for the CD44^high^ subpopulation, CD44^low^ subpopulation, and unsorted population, respectively (Kruskal-Wallis rank sum test, *P* < 2.2 × 10^−308^). With both batch effects and biological difference among cell populations, LCA inferred five clusters, with cells in Batch 1 being represented by Clusters 1 and 2 and cells in Batch 2 being grouped in Clusters 3 to 5 (**Supplementary Fig. 1D**). Similarly, SC3 inferred a 12-cluster structure. Consistent with the strong batch effects observed in bulk RNA-seq analysis, when required to infer a two-cluster model, both LCA and SC3 achieved a perfect separation based on the batch information (NMI = 1 and 0.99, respectively). Next, we evaluated the association between the batch covariate and individual LC states. When controlled at FDR ≤ 0.05, the first LC state was the only one significantly associated with the batch information (*P* < 2.2 × 10^−308^) (**Supplementary Fig. 1E**). After excluding this state, LCA retrieved an optimal structure with two clusters. Of 7005 cells profiled in the sorted CD44^low^ subpopulation, 6733 (96.1%) were clustered into Cluster 2, suggesting that the subpopulation was relatively pure. However, 1796 (26.6%) of 6757 of cells profiled in the sorted CD44^high^ subpopulation were grouped in Cluster 2 with the CD44^low^ cells (**Fig. 5A**). We evaluated the expression signatures for DEGs identified in bulk RNA-seq (log2FC ≥ 1, FDR ≤ 0.05, FPKM ≥ 1 in at least one subpopulation). Of 1709 DEGs (**Supplementary Table 2**), 356 were captured in at least 10% of the cells in one sequenced population or inferred cluster. As expected, Clusters 1 and 2 had higher average expression levels of genes overexpressed in the bulk-sorted CD44^high^ and CD44^low^ subpopulations, respectively (**Fig. 5B**). Importantly, Cluster 2 cells from the sorted CD44^high^ subpopulation had significantly lower expression of those DEGs overexpressed in the bulk CD44^high^ subpopulation than did Cluster 1 cells from the sorted CD44^high^ subpopulation (*P* < 2.2 × 10^−308^) or the sorted CD44^low^ subpopulation (*P* = 7.5 × 10^−71^). Analysis of DEGs overexpressed in the bulk CD44^low^ subpopulation showed the same pattern (**Fig. 5C)**. These results suggested that Cluster 2 cells in the sorted CD44^high^ subpopulation more closely resembled CD44^low^ cells. Moreover, the differential expression pattern was essentially captured by the first remaining LC state (Spearman correlation coefficient = -0.90, *P* < 2.2 × 10^−308^), confirming the biological importance of the inferred LC states. Of the three established molecular markers (*TFAP2B*, *MYOG,* and *NOS1*) for FP-ARMS[21], *TFAP2B* and *MYOG* were overexpressed in the sorted CD44^low^ subpopulation (**Supplementary Table 2**). Moreover, known *PAX3-FOXO1* target genes were significantly enriched in DEGs overexpressed in the CD44^low^ subpopulation (**Supplementary Table 3**). These results suggest that the CD44^high^ subpopulation represents a less-differentiated, stem-like cell subpopulation. Although the exact mechanism by which the distinct subpopulations develop warrants further investigation, we conclude that LCA can control technical variations and reveal reliable transcriptome-based heterogeneity.

**figure 5.**
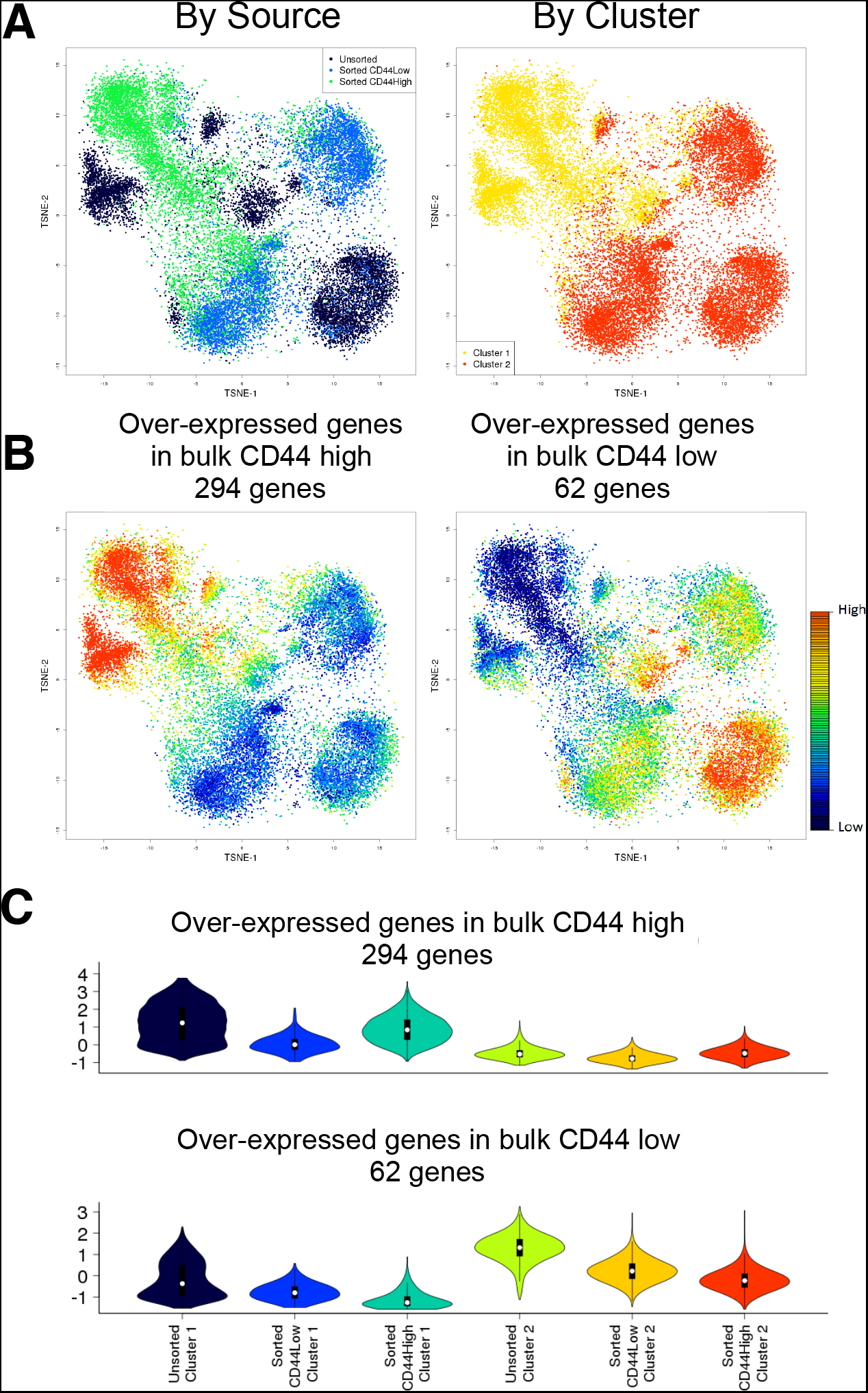
LCA analysis of Rh41 cells, correcting for batch effects. (A) t-SNE plots of clustering results, colored according to source ID or cluster ID. (B) Cellular expression patterns of DEGs identified in bulk RNA-seq. (C) Violin plot of expression patterns of DEGs from bulk RNA-seq in cells classified by source and cluster ID.

## Discussion and conclusions

The rapid technological advance in scRNA-seq platforms has inspired a tremendous explosion of data-analysis methods for identifying heterogeneous subpopulations. Most methods employ a gene-selection step before clustering analysis, based on the assumption that a small subset of highly variable genes is useful for revealing cellular diversity. Although this assumption is valid in most scenarios and reduces the data dimensionality, it potentially excludes genes that are informative for separating subpopulations with subtle diversity. Also, in datasets with strong batch effects, it may result in a small set of retained genes being dominated by batch effects. Moreover, several methods require the user to provide an estimation of the number of clusters in the data, and this may not be readily available.

Using a dual-space search strategy, LCA bypasses the gene selection, learns biologically informative cellular states directly from the raw gene expression matrix, reduces potential technical variations, and measures the cell-to-cell distance by using cosine similarity in the low-dimensional and informative cellular-state space in a data-driven and unsupervised fashion. Cosine similarity has been widely used in information retrieval and text mining to reveal the relation between text documents, a process that shares many similarities with scRNA-seq analysis [22]. Furthermore, LCA provides a mathematical solution for assigning new cells to inferred clusters in a model learned from a subset of cells, a capability that is urgently needed to handle the ever-increasing sample sizes in scRNA-seq. We have demonstrated through extensive simulation and the use of large-scale scRNA-seq datasets that LCA is an efficient, scalable, and robust clustering algorithm that outperforms other tools without the explicit need for gene selection or an estimation of the number of clusters in the data.

## Methods

### Latent cellular states

The input to LCA is a gene expression matrix in which each column is a cell, and each row is a gene/transcript. In UMI-based platforms, the expression level of a gene in a cell is divided by the total expression in that cell to generate a relative expression matrix (T). In read-count based platforms, T can be derived from size factor normalized expression measures. The relative expression matrix is then log-transformed after adding a zero-correction term:

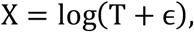

where *∊* is an arbitrarily small number.

We obtain the LC states from a singular value decomposition (SVD) of **X**.

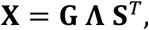

where **S** is a cell-by-LC states matrix. We note that under certain conditions, **S** is the same as the loading matrix from the principle component analysis (PCA) result of **X** [23, 24].

### Determination of significant LC states

We apply the Tracy-Widom test to associated eigenvalues to determine which LC states are significant [10, 25, 26]. The LC state associated with the eigenvalue *λ* is significant if it is significantly different (*P* < 0.05) from the Tracy-Widom distribution, with

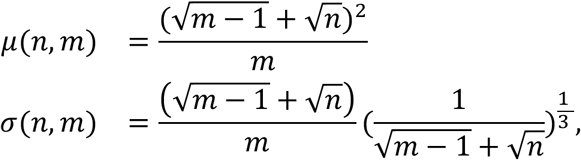

where *n* is the total number of genes and *m* is the total number of LC states. We then discard all the LC states that are not significant or strongly associated with known technical variations (e.g., cell-specific library complexity and batch information), leading to a lower-dimensional cell–by–LC states matrix **S**.

### Distance calculation

Distances between cells in **S** (the cell–by–LC states matrix) are calculated using cosine distance:

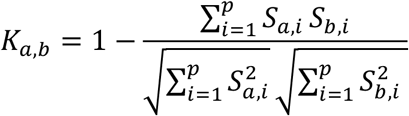

where *S_ai_* and *S_bi_* represent LC state *i* for cells *a* and *b*, and *p* is the total number of retained LC states.

### Spectral clustering

We perform spectral clustering [27] on the resulted distance matrix to derive a set of candidate clustering models with a range of cluster numbers (i.e., 2 – 20 by default).

### Distance measure in the PC space

The cell-gene relative expression matrix was decomposed by PCA and the significant components were determined using the Tracy-Widom test. Distance between cells was measured by the correlation distance of significant components.

### The optimal number of clusters and informative cellular states

We rank the candidate clustering solutions (with different number of clusters) by the silhouette score [11] measured in the PC space. With 2 or more clusters, the silhouette measures the similarity of an individual to its cluster, as compared to other clusters. For each cell, let *d_b_* be the lowest dissimilarity to any other cluster and let *d_w_* be the average dissimilarity to other cells in its cluster. We calculate the silhouette as

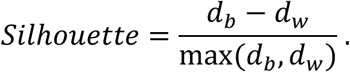

We assign a silhouette score of zero for the default solution of one cluster. An end-user may evaluate the top candidate solutions to determine the optimal number of solutions or specify it with a priori biological knowledge.

With the selected number of clusters, we retain LC states that show significant difference among candidate clusters, update the distance matrix and derive the final clustering solution.

### Fast processing of large numbers of cells

We classify unknown cells **X**′ efficiently by projecting them into space spanned by the informative LC states learned from a representative sample:

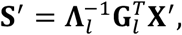

where **Λ***_l_* is calculated from the original **Λ** by removing those rows and columns that are not associated with the informative LC states, and **G***_l_* is calculated from **G** by removing those columns that are not associated with the informative LC states. We can then calculate the cosine similarity between **S**′ and an “average” cell from each cluster and find the cluster with the maximum similarity for each unknown cell.

### Simulation of different cell types

We simulated different cell types from a relatively homogeneous population of cells. We started from the scRNA-seq dataset for sorted CD44^high^ Rh41 cells. The data represented 6757 cells with 20,709 genes detected, with 86% of the entries in the data matrix being zeroes. We considered two factors that have substantial effects on the performance of clustering analysis: the sample size and the degree of difference between cell types.

We tested five different sample sizes (the total number of cells analyzed): 250, 500, 1000, 2000, and 4000 cells. We sampled the selected number of cells from the original set of cells then randomly partitioned them into three sets of variable size, ranging between 10% and 50% of the total number of cells.

To test the effect of the degree of difference between cell types, we varied the number of DEGs and the simulated difference in gene expression. We chose three sets of genes such that the average expression levels for each set were *𝔤*_1_, *𝔤*_2_, and *𝔤*_3_, with 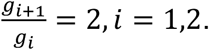 We then rotated the labels of the three sets of genes to make three different sets of cells with perturbed expression: *𝔤*_1_, *𝔤*_2_, *𝔤*_3_; *𝔤*_2;_, *𝔤*_3_, *𝔤*_1_; and *𝔤*_3_, *𝔤*_1_, *𝔤*_2_. We tested two different numbers of DEGs: 300 and 1200. The simulation setting is summarized in **Supplemental Table 4**. For each set of parameters, we generated 100 random samples for testing.

### Normalized mutual information and variation of information

We evaluate the performance of clustering against true labels of cells by using normalized mutual information (NMI) [14] and variation of information (VI) [13]. We use the R package *igraph* to calculate both metrics [28].

### Benchmarking

We used SC3 (Version 1.4.2), Seurat (Version 2.0.0), Monocle2 (Version 2.4.0) and SIMLR (Version 1.2.1) for comparisons with LCA. We used the default settings for all the packages tested. For SIMLR, when the size of the sample was 1000 or larger, we used SIMLR_Large_Scale() instead of SIMLR().

### Rh41 single-cell dataset

The human alveolar rhabdomyosarcoma cell line Rh41 was grown in culture in a 5% CO2 incubator in 75-cm^2^ vented flasks containing DMEM medium supplemented with 10% FBS and 2× glutamine until the cells reached 75% confluence at approximately 3.6 × 10^6^ cells. The cells were detached from the flask with 7 mL of 1× citrate saline to which 7 mL of DPBS was added. The cell suspension was then centrifuged at 300 × *𝔤* for 7 min, and the cell pellet was resuspended in 300 μL of blocking buffer (rat IgG/PBS) and incubated on ice for 30 min. An aliquot of 50 μL of the cells in blocking buffer was transferred to a separate tube for the isotype control. The cells were washed with 1 mL of staining buffer (5% BSA/PBS) and centrifuged at 300 × *𝔤* for 5 min. The pellet, which contained approximately 3 × 10^6^ cells, was then incubated with rat IgG2B anti-CD44-Alexa Fluor 488 antibody (R&D Systems) in staining buffer (15 μL antibody + 135 μL of staining buffer) on ice for 30 min. For the isotype control, approximately 600,000 cells were incubated with 5 μL of rat IgG2B–Alexa Fluor 488 (R&D Systems, Minneapolis, MN) + 45 μL of staining buffer on ice for 30 min. After the incubation, both sets of cells were collected by centrifugation, washed with 1 mL of staining buffer as described above, and resuspended in staining buffer. Flow cytometric analysis was then used to identify the fractions corresponding to the CD44 ^high^ and CD44 ^low^ populations.

For the single-cell experiment, Rh41 cells were grown in culture, harvested, and washed in DPBS as described above. They were then resuspended in PBS/0.2% BSA at a concentration of 1 × 10^6^ cells/mL. The 10x Genomics Single Cell platform performs 3′ gene expression profiling by poly-A selection of mRNA within a single cell, which uses a cell barcode and UMIs for each transcript. Single-cell suspensions were loaded onto the Chromium Controller according to their respective cell counts to generate approximately 6000 partitioned single-cell GEMs (Gel Bead-in-EMulsions). The library was prepared using the Chromium Single Cell 3′ v2 Library and Gel Bead Kit (10x Genomics) in accordance with the manufacturer's protocol. The cDNA content of each sample after cDNA amplification for 12 cycles was quantified, and the quality was checked by High-Sensitivity DNA chip analysis on an Agilent 2100 Bioanalyzer (Agilent Technologies, Santa Clara, CA) at a dilution of 1:6. This quantification was used to determine the final library amplification cycles in the protocol, which were calculated out to 12 cycles. After library quantification and a quality check by DNA 1000 chip (Agilent Technologies), samples were diluted to 3.5 nM for loading onto the HiSeq 4000 sequencer (Illumina) with a 2×75-bp paired-end kit, using the following read length: 26 bp Read1 (10x cell barcode and UMI), 8 bp i7 Index (sample index), and 98 bp Read2 (insert). In total, 518 million, 237 million, and 154 million reads were obtained for unsorted, CD44^low^, and CD44^high^ populations, respectively. The Cell Ranger 2.0.1 Single-Cell Software Suite (10x Genomics) was implemented to process the raw sequencing data from the Illumina HiSeq run. This pipeline performed de-multiplexing, alignment (GRCh38/STAR), and barcode processing to generate gene-cell matrices used for downstream analysis.

After matrix generation, the ribosomal and mitochondria-related genes were filtered out.

### Rh41 bulk RNA-seq dataset

RNA was isolated from the sorted subpopulations by using Trizol (Thermo Fisher Scientific) in accordance with the manufacture's recommendations. RNA libraries were prepared with the KAPA RNA HyperPrep Kit with RiboErase (Roche), using the recommended conditions. Briefly, 200 ng of total RNA was used as input for fragmentation, reverse transcription, and second-strand synthesis. After clean-up, end repair, and A-tailing, NEXTflex adapters (Bioo Scientific, Austin, TX) were ligated to the fragments, and this was followed by 12 cycles of PCR amplification on a C1000 Thermal Cycler (Bio-Rad). Paired-end sequencing was performed (151 bases per read) on a HiSeq 4000 system (Illumina). Three replicates were generated. HTSeq [29] was used to produce the count data, and edgeR [30] was used for the DE analysis with TMM normalization. Each replicate was coded as a pair of CD44^high^ and CD44^low^ in the analysis.

### Statistical analysis

If not specifically stated, the statistical significance was evaluated by the Wilcoxon rank sum test.

## Abbreviations

CD: Cluster of differentiation
DEG: Differentially expressed genes
ERMS: fusion-negative embryonal rhabdomyosarcoma
FP-ARMS: fusion– positive alveolar rhabdomyosarcoma
LCA: Latent cellular analysis
NMI: Normalized mutual information
PBMC: Peripheral blood mononuclear cell
PCA: Principal component analysis
t-SNE: t-Distributed stochastic neighbor embedding
UMI: Unique molecular identifier
VI: Variation of Information

## Declarations

### Acknowledgements

We thank Keith A. Laycock for editing the manuscript.

### Availability of data and materials

The Rh41 scRNA-seq dataset generated during the current study is available from the corresponding author upon reasonable request. The bulk RNA-seq data for sorted CD44^high^ and CD44^low^ subpopulations have been deposited in the European Bioinformatics Institute (EMBL-EBI) under accession number EGAS00001002868. The functions used for the data analysis are included in the single_cell_LCA package, which can be installed from Bitbucket (https://bitbucket.org/changdecheng/singlecelllca).

---

**Supplemental Figure 1.**
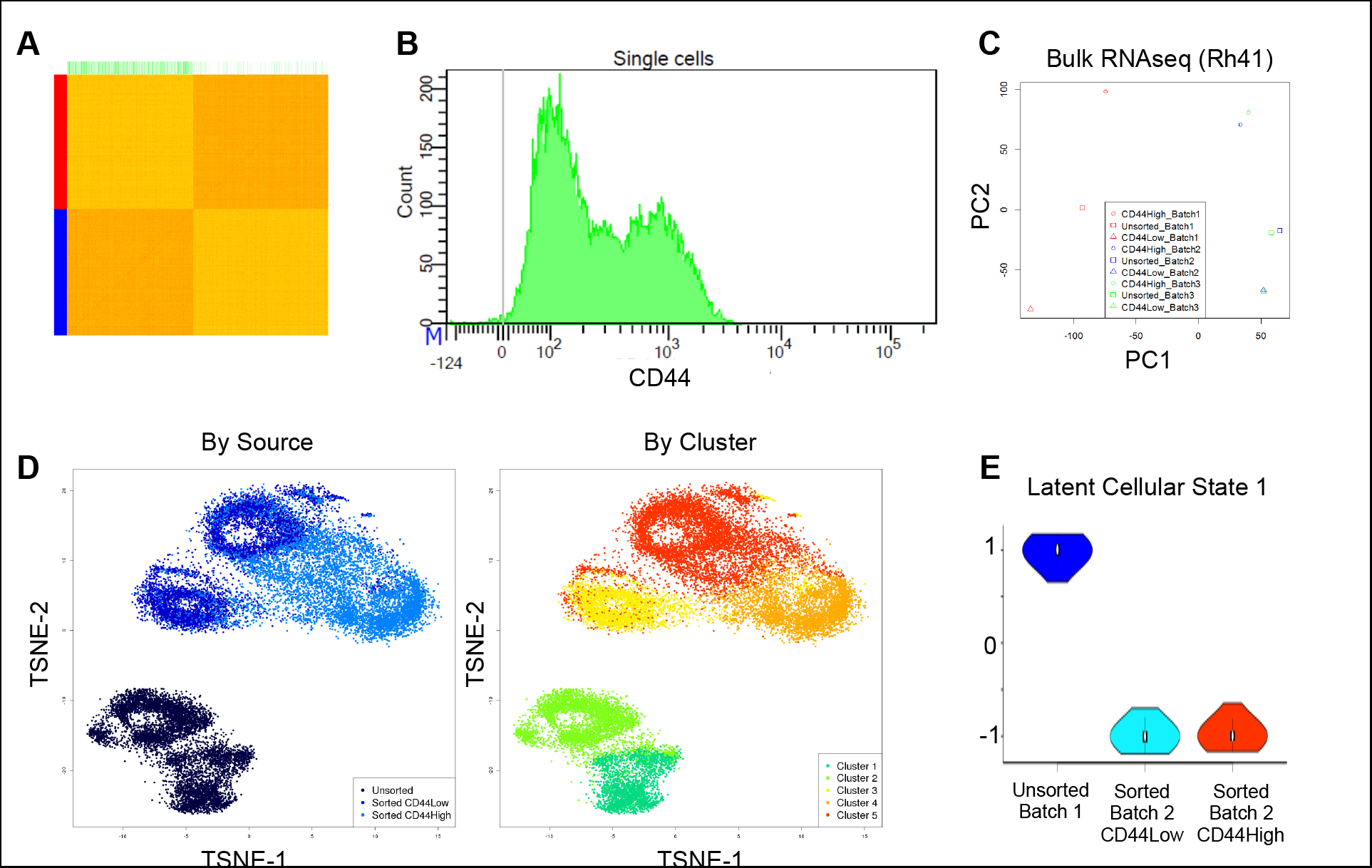
Single-cell analysis of Rh41 cells with strong batch effects. (A) Distance matrix visualized as a heat map. (B) Flow cytometry confirmed a bimodal expression pattern of CD44 in Rh41 cells. (C) The first two PCs for the bulk RNA-seq data showed strong batch effects. (D) t-SNE plot of cells, including sorted and unsorted samples, colored according to source/cluster ID. (E) The first LC state was significantly associated with batch information.

**Supplementary Table 1.** The results of applying LCA and SC3 to the dataset of Tirosh et al.^26^, using normalized mutual information.

|  | LCA | SC3 |
| --- | --- | --- |
| All cells | 0.77 | 0.69 |
| Malignant cells | 0.99 | 0.99 |
| All stromal cells | 0.53 | 0.42 |
| All cells excluding T cells | 0.97 | 0.92 |
| Stromal excluding T cells | 0.90 | 0.71 |

**Supplementary Table 2.** Expression profile of differential expressed genes (in a standalone excel file).

**Supplementary Table 3.** ChEA analysis by Enrichr[31] of 361 overexpressed genes in the CD44^low^ subpopulation of Rh41 (in a standalone excel file).

**Supplementary Table 4.**
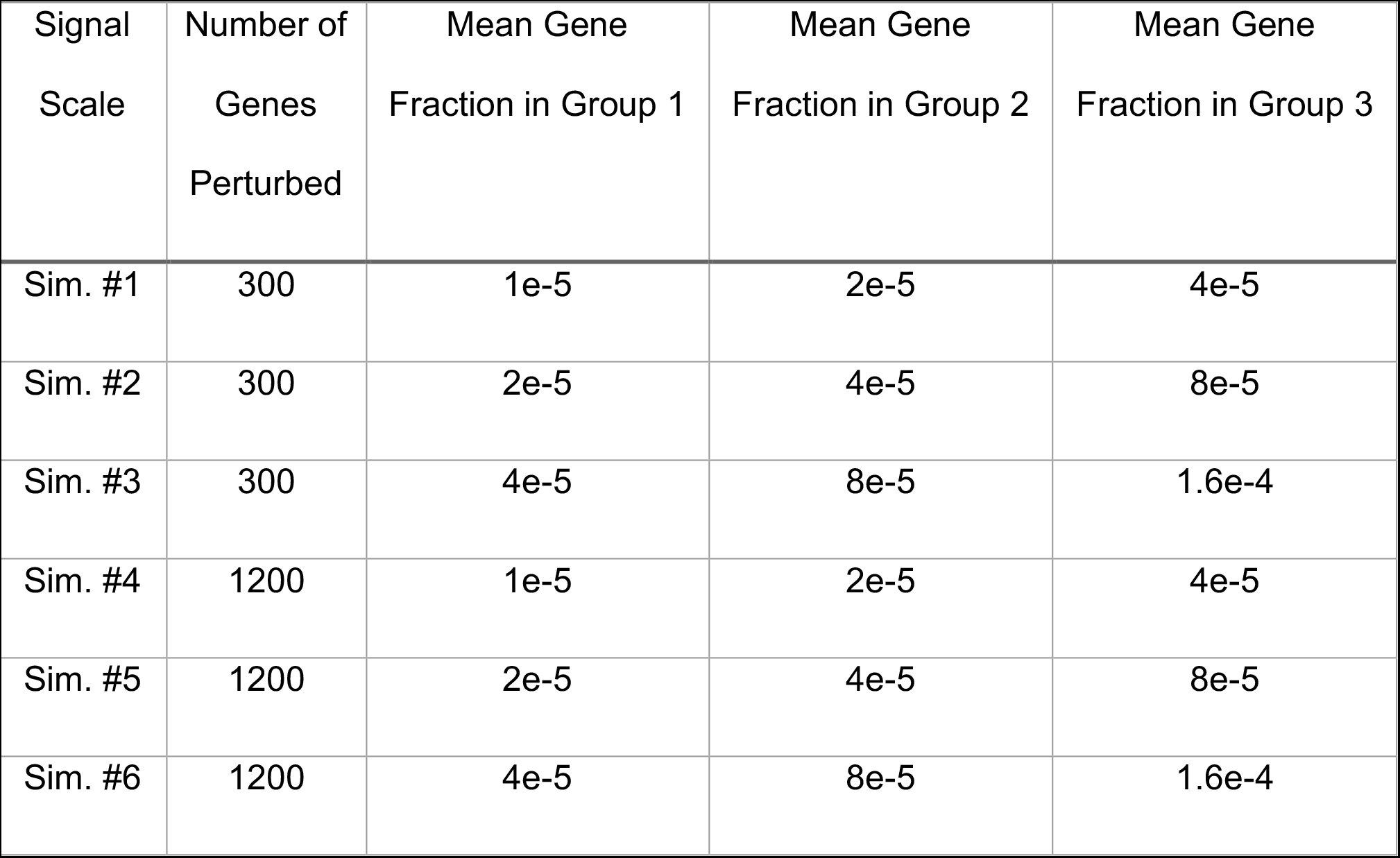
Summary of simulated datasets.

| Signal<br>Scale | Number of<br>Genes<br>Perturbed | Mean Gene<br>Fraction in Group 1 | Mean Gene<br>Fraction in Group 2 | Mean Gene<br>Fraction in Group 3 |
| --- | --- | --- | --- | --- |
| Sim. #1 | 300 | 1e-5 | 2e-5 | 4e-5 |
| Sim. #2 | 300 | 2e-5 | 4e-5 | 8e-5 |
| Sim. #3 | 300 | 4e-5 | 8e-5 | 1.6e-4 |
| Sim. #4 | 1200 | 1e-5 | 2e-5 | 4e-5 |
| Sim. #5 | 1200 | 2e-5 | 4e-5 | 8e-5 |
| Sim. #6 | 1200 | 4e-5 | 8e-5 | 1.6e-4 |

